# Resolving the temporal splenic proteome during fungal infection for discovery of putative dual perspective biomarker signatures

**DOI:** 10.1101/2023.04.08.535756

**Authors:** Benjamin Muselius, Florence Roux-Dalvai, Arnaud Droit, Jennifer Geddes-McAlister

## Abstract

Fungal pathogens are emerging threats to global health with the rise of incidence associated with climate change and increased geographical distribution; factors also influencing host susceptibility to infection. Accurate detection and diagnosis of fungal infections is paramount to offer rapid and effective therapeutic options. For improved diagnostics, the discovery and development of protein biomarkers presents a promising avenue; however, this approach requires a *priori* knowledge of infection hallmarks. To uncover putative novel biomarkers of disease, profiling of the host immune response and pathogen virulence factor production is indispensable. In this study, we use mass spectrometry-based proteomics to resolve the temporal proteome of *Cryptococcus neoformans* infection of the spleen following a murine model of infection. Dual perspective proteome profiling defines global remodeling of the host over a time course of infection, confirming activation of immune associated proteins in response to fungal invasion. Conversely, pathogen proteomes detect well-characterized *C. neoformans* virulence determinants, along with novel mapped patterns of pathogenesis during the progression of disease. Together, our innovative systematic approach confirms immune protection against fungal pathogens and explores the discovery of putative biomarker signatures from complementary biological systems to monitor the presence and progression of cryptococcal disease.

## 1. Introduction

Applications of mass spectrometry-based proteomics for investigating infectious diseases have progressed substantially over the past two decades. With technological improvements and bioinformatic advancement, mass spectrometers provide accurate, cost-effective, and relatively rapid identification of clinical specimens to support diagnostics but with limitations of specificity and speed^1, 2^. Alternative approaches combining liquid chromatography with high-sensitivity and high-resolution tandem mass spectrometry (LC-MS/MS) provides improved identification specificity without the need for microbial culturing to reduce diagnostic processing times^3, 4^. Moreover, combining LC-MS/MS with artificial intelligence (e.g., machine learning, deep learning) algorithms supports species prediction in record time^5^. Beyond identification, mass spectrometry-based proteomics also supports assay development, including biomarkers, to detect pathogen presence within a biological sample and enumerate abundance of the disease-causing agent^6, 7^. Biomarkers are typically designed to detect and quantify changes in the host during disease or to recognize and define a pathogen causing disease^8, 9^. However, given the interplay between a host and pathogen during infection, profiling of biomarkers from both perspectives can be beneficial for diagnosing, monitoring, and treating disease.

For fungal pathogens, the emergence of new species and strains, increased geographical and host ranges, as well as rising rates of antifungal resistance clearly emphasize the need for improved and effective diagnostic and therapeutic options^10, 11^. For the opportunistic human fungal pathogen, *Cryptococcus neoformans,* which infects over 220,000 people/year with mortality rates exceeding 80%^12^ in difficult to treat or untreated infections, novel detection and treatment options are needed^13^. *C. neoformans* is found ubiquitously within the environment, surviving in desiccated plant matter and bird guano, where disruption of these reservoirs promotes aerosolization of the fungus and inhalation by the host supports colonization of the respiratory tract^14^. If the resulting pulmonary fungal infection cannot be cleared by resident macrophages within the lungs, cryptococcosis (e.g., fungal meningitis) may develop following fungal dissemination across the blood brain barrier and transmission to the central nervous system^15^. A key determinant of the persistence and severity of cryptococcal infection is underscored by immunocompetency of the host and virulence factor production by the pathogen. From the host perspective, phagocytosis by M1 differentiated alveolar macrophages, antigen presentation, and recruitment of T cells support an effective response to *C. neoformans*^16^. Whereas, the pathogen will produce a plethora of virulence factors, including a polysaccharide capsule, melanin, and extracellular enzymes (e.g., urease, superoxide dismutase), along with thermotolerance to withstand the host immune response^17, 18^.

The spleen, a secondary lymphoid organ, plays a pivotal role in the timely recognition of a pathogen associated with the innate immune response followed by effective pathogen clearance upon initiation of the adaptive immune system^19^. Composition of the spleen includes white and red pulp that are separated by the marginal zone, which contains highly phagocytic macrophages that support detection and presentation of antigens, such as polysaccharides, from invading pathogens^20, 21^. In this study, we establish a murine inhalation model of cryptococcosis, followed by collection of the spleen and enumeration of fungal burden. Using a systematic approach, we integrate mass spectrometry-based proteomics profiling with tandem mass tags for quantification and multiplexing to reveal temporal axes of both biological systems influenced during infection. We observe tailored responses by the host during an active immune response to defend against infection and reciprocal virulence factor production by the pathogen to sustain the infection. Moreover, we define mapped patterns of fungal virulence that suggest novel associated roles for *C. neoformans* proteins. Finally, by focusing on a critical immune-associated organ (i.e., spleen) that plays important roles in filtering pathogens from the blood during infection, we propose putative protein biomarker signatures from both the host and pathogen perspectives to diagnose and monitor disease progression over time.

## 2. Material and Methods

### 2.1. Fungal strains, growth conditions, and media

*Cryptococcus neoformans* strain H99 (wild-type) was used for all experiments. Colonies were maintained on yeast extract peptone dextrose (YPD) media, grown at 30 °C, and stored at 4 °C unless otherwise specified.

### 2.2. Murine model and tissue collection

All animal utilization protocols were performed under the guidelines and approval of the University of Guelph (AUP #4193). A 5 mL liquid culture of *C. neoformans* in YPD was grown overnight at 30 °C with shaking at 200 rpm and sub-cultured 1:100 into fresh YPD. The sub-culture was grown for 12 h under the same conditions. Cells were washed twice with PBS and cell density was determined with a hemocytometer. Cells were diluted to 4e6/mL in sterile PBS to construct the inoculum. BALB/c mice (60 females) were used in this trial, 30 designated as infected and 30 as uninfected controls. Notably, female mice were selected for this experiment based on preliminary findings of response to infection (data not shown). Control mice underwent all the same procedures as infected mice but received only sterile PBS instead of inoculum. Mice were housed in cages of either three or four individuals/cage with only individuals belonging to the same treatment groups were co-housed. Inoculation was performed under isoflurane induced anesthesia. Mice were brought to surgical plane and the inoculum was administered intranasally, mice were held vertically for ∼1 min post inoculation to ensure no inoculum was lost to regurgitation^22^. Mice were then weighed and placed in an empty cage to recover before being returned to the home cage. Mice were assessed at least once every 24 h on five criteria; respirator difficulty, coat condition, posture, activity level, and weight. Any mouse with >20% weight loss or showing severe decline in one or more of the categories was euthanized immediately. Otherwise, infected mice were sacrificed at their designated time point (3-, 10-, or 21-days post inoculation [dpi]) alongside an uninfected partner. Euthanasia was performed using CO_2_ under anesthesia by isoflurane. Post euthanasia an incision was made along the ventral midline of the thorax and abdomen, the spleen was excised, collected, and flash frozen with liquid nitrogen.

### 2.3. Colony Forming Unit Counts

Spleen samples were weighed and homogenized in 1 mL of sterile PBS using a bullet blender (medium speed for 5 min, repeated twice or until the sample was visually homogeneous). A serial dilution was performed up to 10^−5^ before 100 µL of each dilution was plated on YPD agar with 32 µg/mL chloramphenicol to isolate fungal growth. Plates were incubated for 48 h at 30 °C and colony forming units (CFU) were counted and recorded.

### 2.4. Proteomics sample preparation

Proteomics sample preparation was performed as previously described with modifications^23^. Briefly, tris-HCl (pH 8.5) mixed with a protease inhibitor cocktail tablet was added to splenic samples before the tissue was emulsified using a bullet blender (medium speed for 5 min, repeated twice or until the sample was visually homogeneous). Sodium do-decyl sulphate (SDS; final concentration of 2%) and samples were lysed by probe sonication (30 s on/off; 15 cycles, power 3) followed by addition of dithiothreitol (DTT; 10 mM) and iodoacetamide (IAA; 55 mM). Following an overnight acetone precipitation (80% final) at −20 °C, samples were washed with 80% acetone, dried, and resuspended in 8 M urea/40 mM HEPES, before protein content was assayed by bovine serum albumen (BSA) tryptophan fluorescence assay^24^. Ammonium bicarbonate (ABC) was added at a ratio of 3:1 to sample volume and samples were normalized to 100 µg of protein. Samples were digested with LysC/trypsin at a ratio of 50:1 (protein/enzyme) for ∼18 h, stopped by the addition of 10% (v/v) trifluoroacetic acid, and peptides were purified using C_18_ StAGE (STop And Go Extraction) tips^25^.

### 2.5. Tandem mass tag (TMT) labeling

TMTpro 16plex Label Reagent Set (Fisher Scientific) was used for isobaric labeling as directed by the manufacturer^26^. Briefly, peptides were resuspended in tetraethylammonium bromide (TEAB) and standardized to 10 µg per channel and labeled at a ratio of 1:10 (sample:tag). Two standard channels were included, an organ standard containing equal parts (by concentration) of all spleen samples, and a *C. neoformans* standard containing 10 µg *C. neoformans* proteins prepared as previously described^23^. Excess label was removed by Pierce Peptide Desalting Spin Column (Fisher Scientific) as directed by the manufacturer.

### 2.6. High pH fractionation

Multiplexed TMT samples were fractionated using a Pierce High pH Reversed-Phase Peptide Fractionation Kit (Fisher Scientific) used as directed by the manufacturer. Wash fractions were retained as directed and assayed; no proteins were detected in these fractions.

### 2.7. Mass spectrometry

Each vacuum-dried peptide fraction was resuspended and diluted at an estimated concentration of 0.05 µg/µL with 0.1% formic acid in water and an equivalent of 1 µg peptide was loaded on an Evotip disposable trap column (Evosep Biosystems) according to the manufacturer protocol prior to mass spectrometry analysis. LC-MS/MS analysis was performed on an Evosep One liquid chromatography system (Evosep Biosystems) interfaced with an Orbitrap Exploris 480 mass spectrometer (Thermo Fisher Scientific). The chromatographic separation was performed using a manufacturer-defined 60 samples per day method allowing peptide elution with a 21 min gradient of 0.1% formic acid in acetonitrile at a flowrate of 1 µL/min on an Evosep EV1106 separation column (8 cm long, 150 µm internal diameter and 1.5 µm beads). The mass spectrometer was operating in Data Dependent Acquisition in positive mode using the Thermo XCalibur software version 4.6.67. Full scan mass spectra (400 to 1400 *m/z*) were acquired at a resolution of 120,000 with a normalized AGC of 100% and a maximum injection of 50 ms. Each MS scan was followed by acquisition of fragmentation MS2 spectra of the most intense ions above a 5×10^3^ intensity threshold for a total cycle time of 1.5 s (top speed mode). The selected ions were isolated in a window of 0.7 *m/z* and fragmented by Higher energy Collision induced Dissociation (HCD) with 32% of normalized collision energy. The resulting fragments were detected in the Orbitrap at a resolution of 45,000, with a normalized AGC of 200% and a maximum injection time of 120 ms. The first mass of the MS2 spectra mass range was set at 110 *m/z* to ensure the detection of all TMT reporter ions. Internal calibration using lock mass on the *m/z* 445.12003 siloxane ion was used and a dynamic exclusion of previously fragmented peptides was set for a period of 45 s and a tolerance of 10 ppm.

### 2.8. Data processing

MaxQuant software (version 2.1.3.0) was used for mass spectrometry raw data file processing^27^. The reference *Mus musculus* (55,260 proteins, downloaded: 21 September 2022) and *C. neoformans* (7,429 proteins, downloaded: 21 September 2022) proteomes retrieved from Uniprot (https://www.uniprot.org) were searched using the inbuilt Andromeda search engine^28^. Trypsin enzyme specificity (two missed cleavages), fixed modifications (carbamidomethylation of cysteine), and variable modifications (methionine oxidation, N-acetylation). Reporter ion was set to MS2, 16plex selected, and correction factors were imported from manufacturer documentation. Normalization was set to weighted ratio to reference channel (i.e., organ standard), isobaric weight exponent was set to 0.75 with re-quantification enabled, a minimum peptide length of seven amino acids, and a minimum to two peptides were required for identification.

### 2.9. Bioinformatics

Bioinformatic processing was performed with Perseus (version 2.0.6.0)^29^. Data was filtered to remove potential contaminants, reverse hits, and proteins identified by site. A log_2_ transform was applied to all intensity values, a valid value filter (6 of 10 replicates) was applied, missing values were imputed from a normal distribution originating at the group mean with a width of 0.3 and downshift of 1.8. Changes in intensity between treatment and time point were evaluated for statistical significance using a Student’s *t*-test and Benjamini-Hochberg false discovery rate (FDR) correction with a cutoff of 0.05 with a minimum fold change of 2 (S_0_=1). Similarity of trends between time points was determined using a Pierce correlation. A 1D annotation enrichment was performed as previously described (testing whether changes in abundance for proteins within a defined category increase or decrease relative to the normal distribution for that category)^30^. For interaction network observations, a STRING analysis was performed (https://string-db.org).

## 3. Results

### 3.1. The proteome remodels by time and infection status

To explore the relationship between the host and pathogen during cryptococcosis, we established a murine model of infection and collected the spleen, a critical organ for host immune response to infection, for quantitative proteome profiling (Figure 1). The time points of 3, 10, and 21 dpi were selected to represent an early, innate-driven response associated with antigen presentation in the spleen (3 dpi), a mid, mounted immune response against the pathogen (10 dpi), and a late, adaptive immune response with the host succumbing to infection (21 dpi). Our global analysis identified 3,119 proteins (after valid value filtering) with 2,373 detected in the host (*Mus musculus*) and 746 from the pathogen (*C. neoformans*) (Figure 2A). A principal component analysis (PCA) of the global response distinguished component 1 (28%) based on time across the experiment (3, 10, 21 dpi) and component 2 (12.3%) was defined by the infection status at 3 dpi, likely due to immediate immune activation within the spleen upon antigen presentation (Figure 2B).

**Figure 1:**
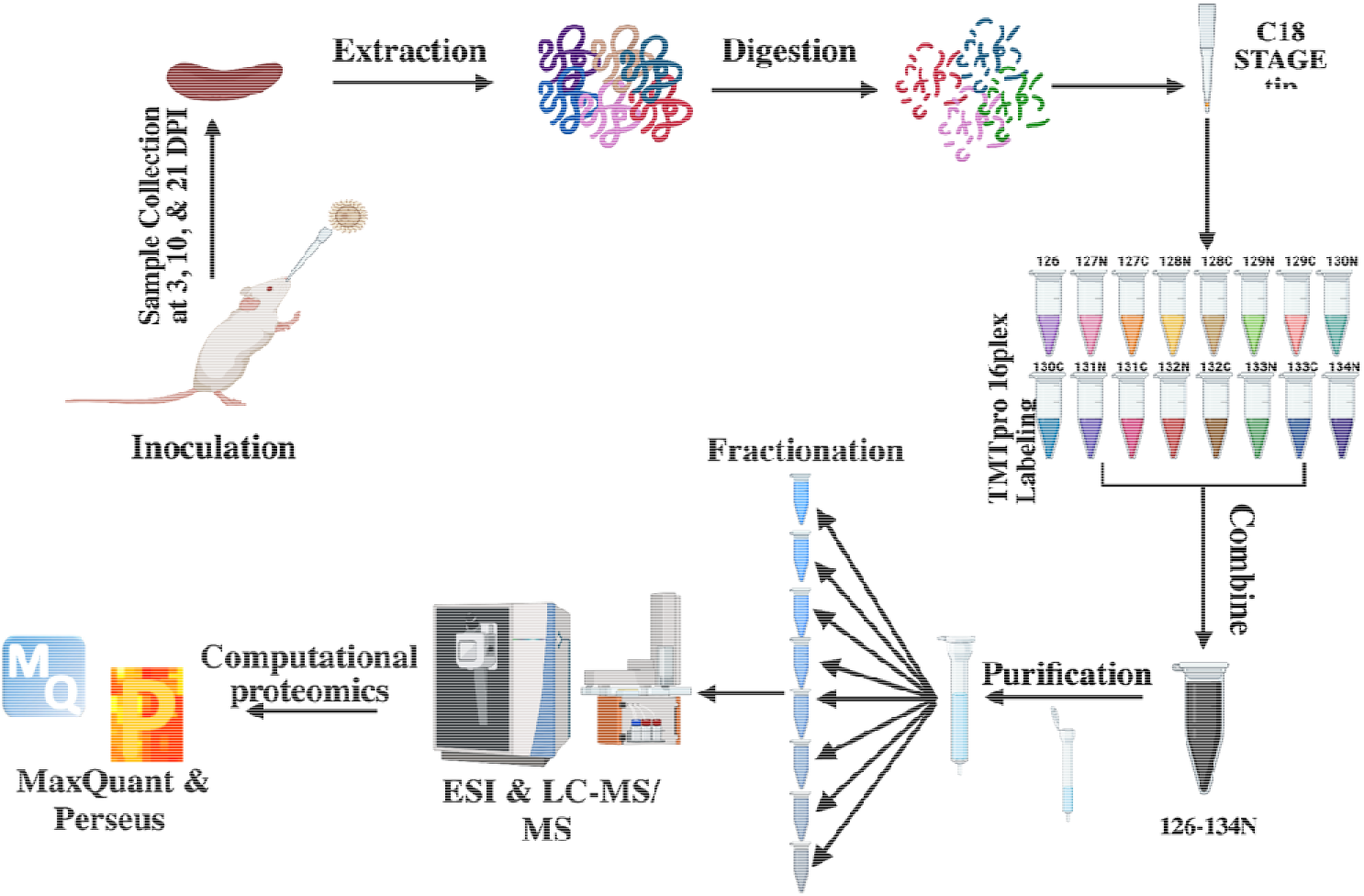
Mass spectrometry-based proteomics workflow. For the *in vivo* cryptococcal infection model, 60 BALB/c female mice (7-9 weeks-old) were divided into uninfected (30 mice) and infected (30 mice) groups and culled at 3, 10, and 21 dpi. The spleen from each mouse was collected and processed with our proteomics protocol. Tandem mass tag labeling was used for multiplexing, samples were fractioned by high pH, and measured on an Exploris240 mass spectrometer. Data analyzed with MaxQuant^27, 29^ and processed with Perseus. Figure generated with Biorender.com.

**Figure 2:**
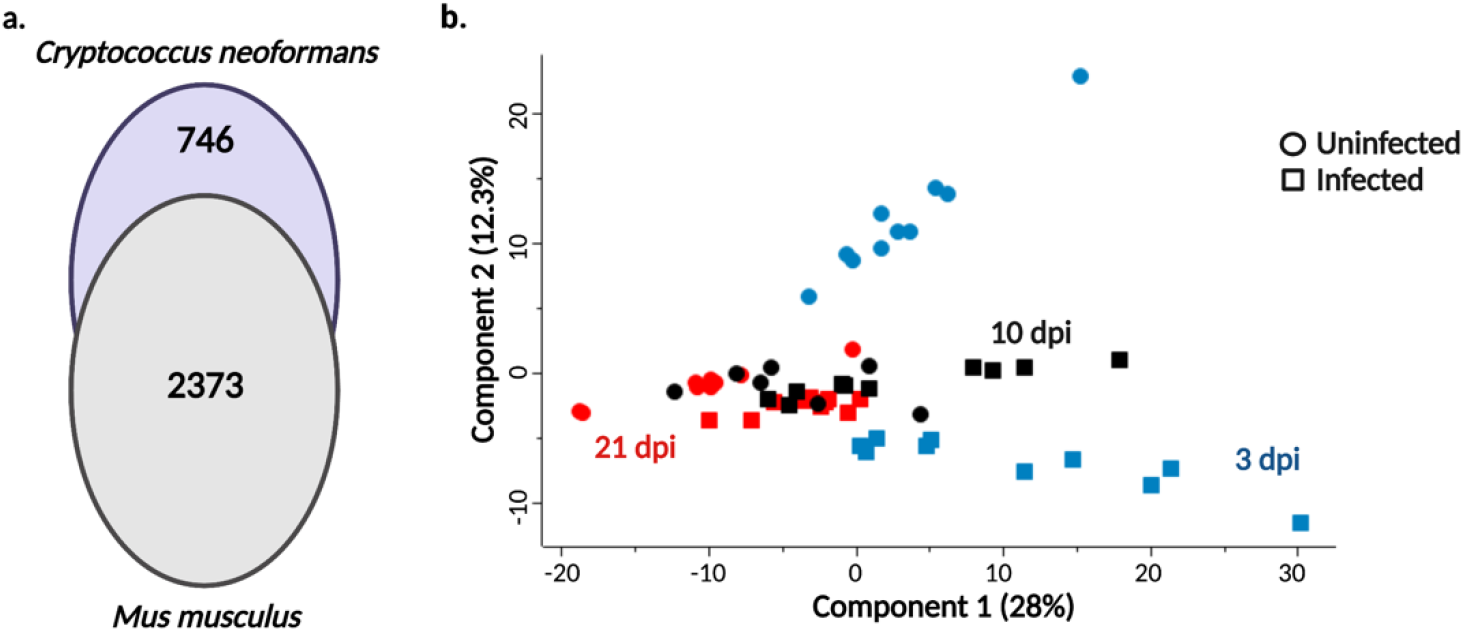
Global proteome overview of temporal cryptococcal infection of the spleen. A) Venn diagram for quantified proteins from spleen for the host (*Mus musculus*) and pathogen (*Cryptococcus neoformans*). B) Principal Component Analysis for infected (squares) vs. uninfected (circles) samples collected at 3 days post inoculation (dpi; blue), 10 dpi (black), and 21 dpi (red). At each time point an uninfected mouse was culled to time match the infected mouse. Experiment performed with 10 biological replicates.

### 3.2. Infection state of the host drives innate and adaptive immune responses

During an infection, the spleen functions to filter the blood of pathogens and abnormal cells, connect antigen presenting cells with T and B cells for immune response activation, recruit immune cells to the sight of infection, and clear the pathogen^19, 31^. Here, we defined a core spleen proteome of 2,233 proteins common between infectious states with the production of 78 proteins regulated by infection and 62 proteins regulated by the absence of infection (Figure 3A). Time was also an important driver of the splenic response with 1,870 proteins produced independent of time, 39 proteins specific to the early responses (3 dpi), 51 proteins unique to the mid response (10 dpi), and 77 proteins produced upon terminal stages of infection (21 dpi) (Figure 3B). A PCA associated the time of infection across both components with component 1 (35.1%) driven by time and component 2 (16.5%) distinguished by immune response diversity at 10 dpi (Figure 3C). To validate activation of the innate and adaptive immune system upon infection, we mapped associated innate (64 proteins) and adaptative (26 proteins) functions for host proteins based on Gene Ontology Biological Processes (GOBP). We observed a significant increase in protein production for designated innate and adaptive proteins under the infected conditions (across all time points) compared to the uninfected samples (Figure 3D). Together, these data confirm immune response activation and demonstrate host proteome remodeling aligned with the infectious state of the host.

**Figure 3:**
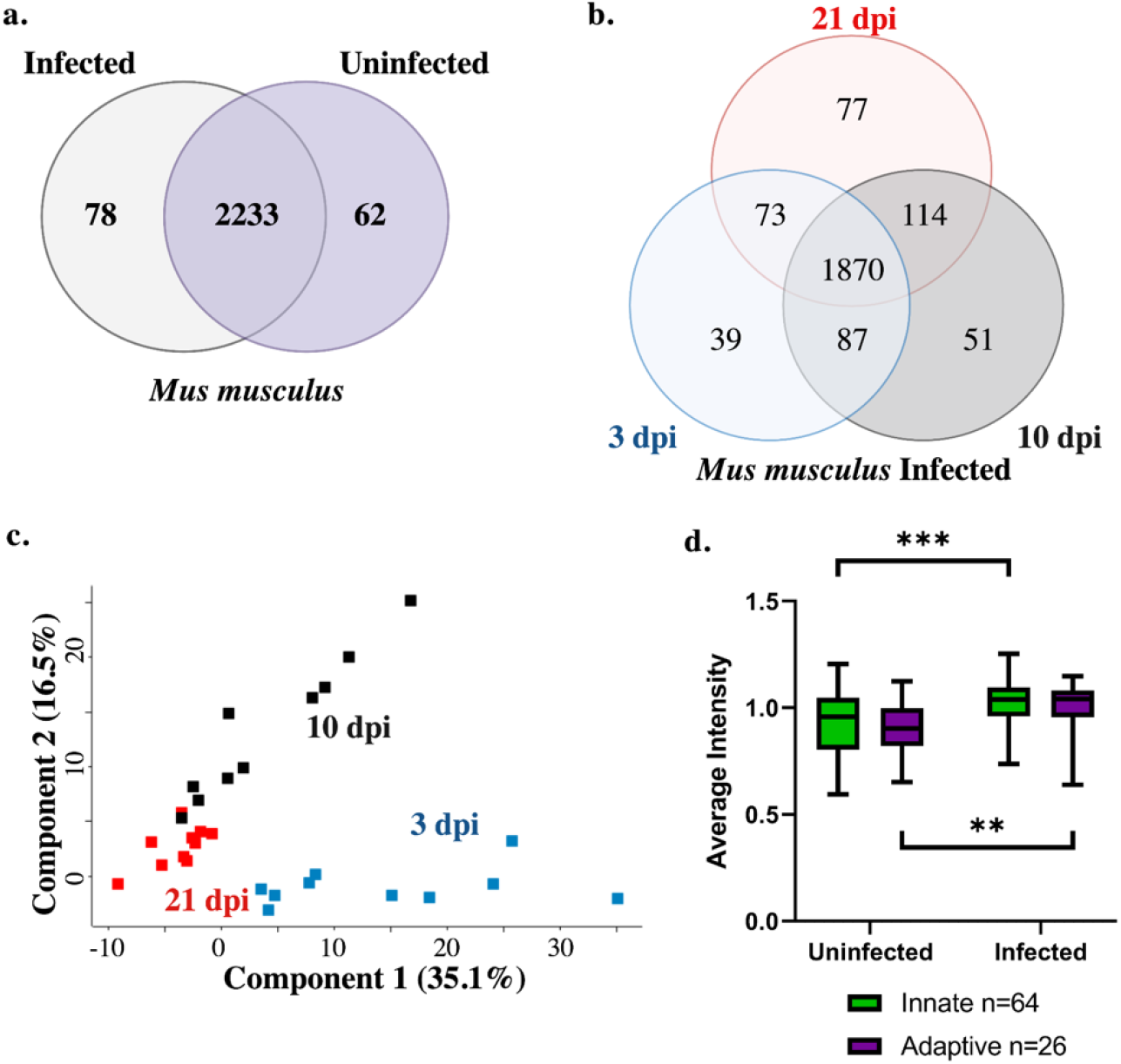
Host-specific response to cryptococcal infection. A) Number of proteins common between uninfected and infected splenic samples (2233 proteins), unique to infected (78 proteins), or specific to uninfected (62 proteins) samples. B) Temporal comparison of identified proteins. A core proteome of 1870 proteins was observed with 39 proteins unique to 3 dpi, 51 proteins unique to 10 dpi, and 77 proteins unique to 21 dpi. C) Principal Component Analysis of host proteins from infected samples over time. D) Average intensity of identified host proteins associated with innate immunity (n = 64) and adaptive immunity (n = 26) based on GOBP across the time points for infected vs. uninfected samples. Welch’s t-test, **p-value < 0.01; ***p-value <0.001.

### 3.3. Anticipated host-specific responses to cryptococcal infection drive temporal changes

Next, we aimed to define significant changes in protein production across the sequential time points and to define common regulators of the temporal immune response. An initial validation of temporal infection processes within the spleen detected the presence of proteins with defined roles in host immune response to fungal pathogens, including activation of lectin-like proteins and the complement cascade^32^. We observed a significant increase in production between 10 and 21 dpi for lectin-like proteins (Figure 4A) and complement-associated proteins (Figure 4B). Given these temporal differences, we performed a comparison between the early- and mid-collection times and identified 230 proteins with a significant increase in abundance at 10 dpi and 142 proteins with a significant increase in abundance at 3 dpi (Figure 4C). A comparison between mid- and late-collection times detected 192 proteins with a significant increase in abundance at 21 dpi and 292 proteins with a significant increase in abundance at 10 dpi (Figure 4D).

**Figure 4:**
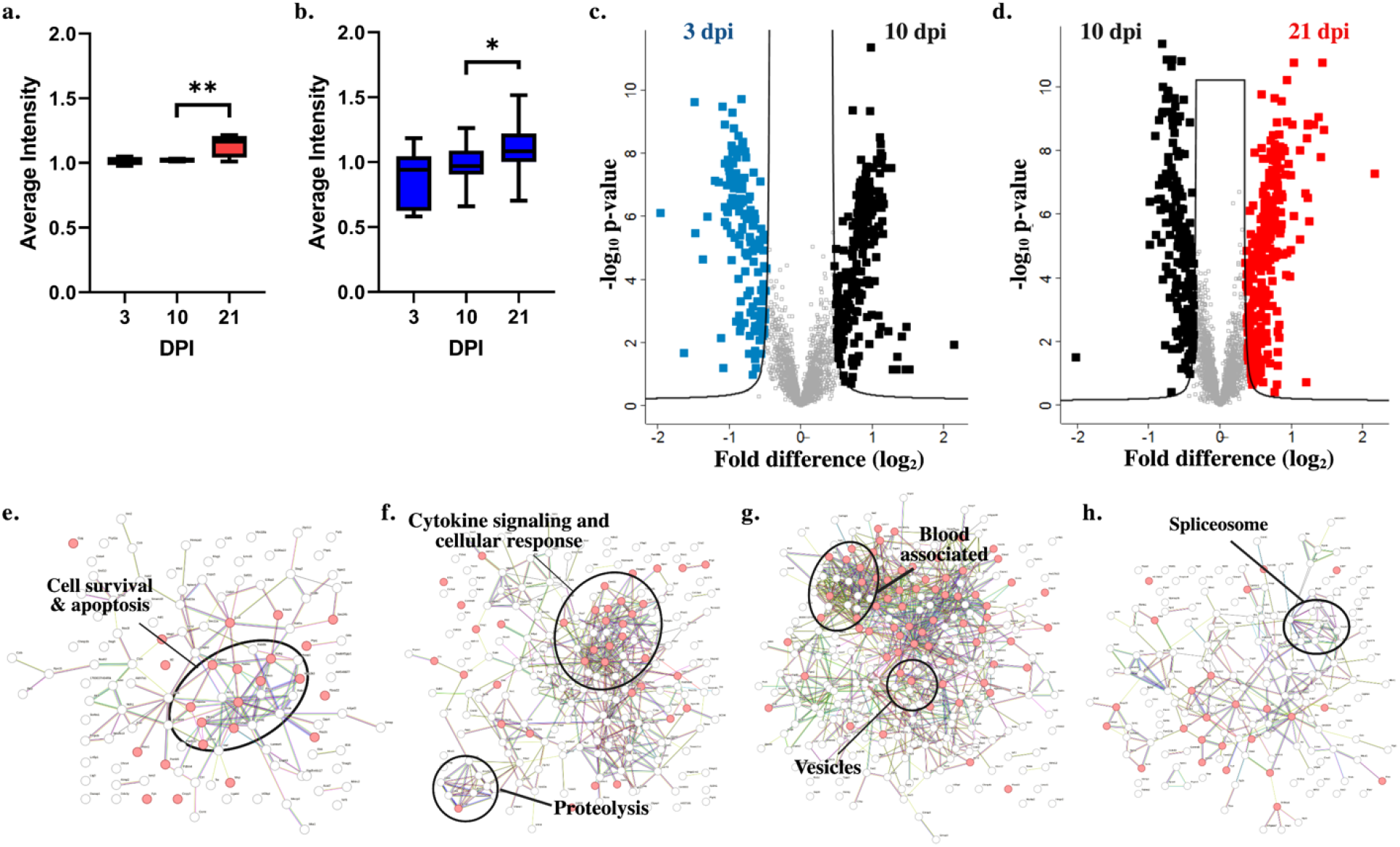
Significant differences in host production demonstrate immune-specific responses. A) Lectin-like proteins by GOBP (n = 4). B) Complement cascade proteins by GOBP (n = 19). C) Volcano plot for significantly different proteins with higher production at 3 dpi (blue) vs. 10 dpi (black). D) Volcano plot for significantly different proteins with higher production at 10 dpi (black) vs. 21 dpi (red). E) STRING interaction map for host proteins with significant increases in abundance at 3 dpi (compared to 10 dpi). F) STRING interaction map for host proteins with significant increases in abundance at 10 dpi (compared to 3 dpi). G) STRING interaction map for host proteins with significant increases in abundance at 10 dpi (compared to 21 dpi). H) STRING interaction map for host proteins with significant increases in abundance at 21 dpi (compared to 10 dpi). For box plots, Welch’s t-test *p-value < 0.05; **p-value < 0.01. For volcano plots, Welch’s t-test p-value < 0.05, FDR = 0.05, S_0_ = 1.

A closer look into the drivers of temporal response by collective functions of proteins with significantly altered abundance profiles based on interaction networks was performed using the STRING database. At 3 dpi, elevated proteins showed clustering associated with host cell survival and apoptosis (Figure 4E) compared to enhancement of proteins involved in cytokine signaling and cellular response as well as proteolysis elevated at 10 dpi (Figure 4F). Conversely, a comparison of 10 dpi to the late time point showed clustering of blood-associated proteins and vesicular functions (Figure 4G). Whereas interactions among the splicesome were observed at 21 dpi (Figure 4H). Together, these data emphasize the critical activation of cytokine signaling as a mode of host defense at 10 dpi to promote a mounted immune response against fungal infection.

### 3.4. Prediction of putative host protein biomarkers based on immune system activation

Given on our observed regulation of the immune response upon infection and throughout the time course of infection, we mapped patterns of protein production for infection trends and classically-defined immune response proteins associated with fungal infections (i.e., complement, lectin-like) (Table 1). For instance, we mapped the host protein, Metaxin-2 (Mtx2; O88441) that regulates Bak, a proapoptotic member of the Bcl-2 family, upon TNF⍰ (tumor necrosis factor-alpha)-induced cell death^33^, to reduced protein production upon infection (Figure 5A). Following this trend, four additional host proteins with mapped patterns of production were identified, including a transcriptional regulator (E9PZM4), hydrolase (Q9CR35), transporter (Q9D3D9), and guanine releasing factor (Q9QUG9). Notably, the hydrolase (chymotrypsinogen B) is a known component of the lysosome with roles in protein digestion and was previously described as a biomarker for cancer^34^. Next, we mapped a host protein with a distinct increase in production upon infection, cathelicidin antimicrobial peptide (Camp; P51437), with demonstrated antifungal properties^35, 36^. Proteins aligning with this candidate include an iron transporter (P08071)^37^, peroxidase (P49290)^38^, antimicrobial (Q61646)^39^, and hormone (Q8K426)^40^. Importantly, each of these mapped proteins has defined roles in host immunity, as well as antimicrobial activity, supporting the validity of our approach to identify relevant host proteins with similar production profiles to key immune regulators.

**Figure 5:**
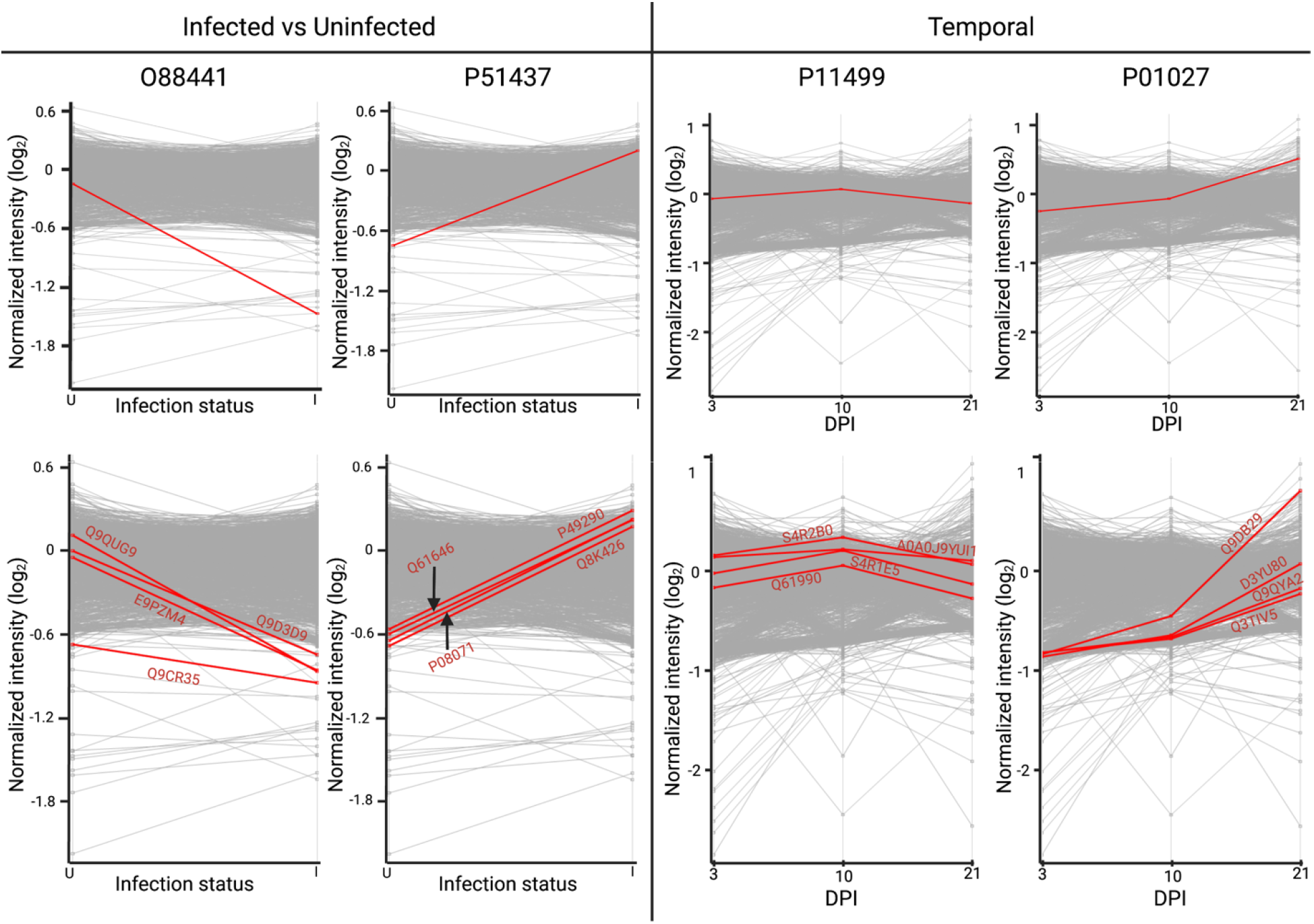
Mapped temporal production for immune-associated host proteins. A) O88441 (Metaxin-2) host protein mapped abundance trend at 3, 10, and 21 dpi. Top four host proteins with comparable mapped protein production based on trend. B) P51437 (Cathelicidin antimicrobial peptide) host protein mapped abundance trend at 3, 10, and 21 dpi. Top four host proteins with comparable mapped protein production based on trend. C) P11499 (Heat shock protein 90-beta) host protein mapped abundance trend at 3, 10, and 21 dpi. Top four host proteins with comparable mapped protein production based on Pearson correlation. D) P01027 (Complement C3) host protein mapped abundance trend at 3, 10, and 21 dpi. Top four host proteins with comparable mapped protein production based on Pearson correlation.

**Table 1:** Putative host protein biomarkers based on mapped patterns of production.

| **Parent Trend Protein** | **Protein Name** | **Gene ID** | **Protein ID** | **Keywords** |
| --- | --- | --- | --- | --- |
| **O88441** | **Metaxin 2** | **Mtx2** | **O88441** | **Protein transport** |
|  | Chromodomain-helicase-DNA-Binding protein 2 | Chd2 | E9PZM4 | Chromatin regulator, Helicase, Hydrolase, Myogenesis, Transcription regulation |
|  | Chymotrypsinogen B | Ctrb1 | Q9CR35 | Hydrolase, Serine protease, Digestion |
|  | ATP synthase subunit delta, mitochondrial | Atp5f1d | Q9D3D9 | ATP synthase, Hydrogen ion transport |
|  | RAS guanyl-releasing protein 2 | Rasgrp2 | Q9QUG9 | Guanine-nucleotide releasing factor |
| **P51437** | **Cathelicidin antimicrobial peptide** | **Camp** | **P51437** | **Antibiotic, Antimicrobial, Innate immunity** |
|  | Lactotransferrin | Ltf | P08071 | Hydrolase, Serine protease, Immunity, Iron transport |
|  | Eosinophil peroxidase | Epx | P49290 | Oxidoreductase, Hydrogen peroxidase |
|  | Haptoglobin | Hp | Q61646 | Antibiotic, Antimicrobial, Antioxidant, Serine protease homolog, Acute phase, Immunity |
|  | Resistin-like gamma | Retnlg | Q8K426 | Hormone |
| **P11499** | **Heat shock protein 90-beta** | **Hsp90ab1** | **P11499** | **Chaperone, Stress response** |
|  | Ubiquitin-conjugating enzyme E2K | Ube2k | A0A0J9YUI1 | ATP-binding, Nucleotide-binding |
|  | Poly(rC)-binding protein 2 | Pcbp2 | Q61990 | DNA-binding, RNA-binding, Ribonucleoprotein, Antiviral defense, Innate immunity |
|  | RNA binding motif protein 25 | Rbm25 | S4R2B0 | RNA-binding |
|  | Glutathione peroxidase | Gpx4 | S4R1E5 | Oxidoreductase, Peroxidase |
| **P01027** | **Complement C3** | **C3** | **P01027** | **Complement pathway, Complement alternate pathway, Immunity, Inflammatory response, Fatty acid metabolism** |
|  | Cold-inducible RNA-binding protein | Cirbp | D3YU80 | RNA-binding, Stress response |
|  | Zinc finger CCCH domain- containing protein 15 | Zc3h15 | Q3TIV5 | Metal-binding, Zinc |
|  | Isoamyl acetate-hydrolyzing<br>esterase 1 homolog | lah1 | Q9DB29 | Hydrolase, Lipid<br>degradation, Lipid<br>metabolism |
|  | Mitochondrial import receptor<br>subunit TOM40 homolog | Tomm40 | Q9QYA2 | Porin, Ion transport,<br>Protein transport |

We then selected temporally-defined mapping profiles for proteins serving validation roles in lectin-like receptors and complement cascade. Here, the heat shock protein 90 ß (Hsp90; P11499), which is a main chaperone protein involved in the activation of the JAK-STAT pathway to promote immune cell differentiation^41, 42^, showed heightened production at 10 dpi compared to 3 and 21 dpi (Figure 5C). Other proteins with mapped patterns of production, including nucleotide binding (A0A0J9YUI1; Q61990; S4R2B0) and a peroxidase (S4R1E5), with the latter protein playing an important role in protection from oxidative damage^43^. For the complement cascade, we mapped C3 (P01027), a central component of the classical and alternative complement pathways involved as an opsonin and for promotion of inflammation, shows elevated production over time (Figure 5D). Additional proteins with similar mapped patterns of production include stress response (D3YU80), metal binding (Q3TIV5), hydrolase (Q9DB29), and protein transport (Q9QYA2) with regulatory roles within the cell. Critically, several of the selected putative host biomarker proteins are involved in immunity and antimicrobial defense with distinct responses over time and infection. However, with ubiquitous roles in host defense, the sole use of these proteins as biomarkers of fungal infection may be limited but adoption of a combination of markers may prove more promising.

### 3.5. The pathogen perspective of infection defines temporal proteome changes associated with increased fungal burden over time

From the pathogen’s perspective, the success of infection defends on many factors, including susceptibility of the host, localization, avoiding immune system recognition, and the production of virulence factors to drive infection forward^44^. Our infectome profiling of the spleen following *C. neoformans* infection showed a consistent core proteome of 578 fungal proteins across the time points with 29 proteins specifically produced at the early infection stage (3 dpi), 20 points unique to the mid-collection time point (10 dpi), and four proteins produced at late-stage infection (21 dpi) (Figure 6A). A PCA defined separation of component 1 (28.8%) by the time course of infection and component 2 (22%) by factors driving distinction between the early and mid-collection times (3 & 10 dpi) vs. the late-stage collection (21 dpi) (Figure 6B). To tease apart potential drivers of the distinction for *C. neoformans* protein production at 21 dpi, we performed a fungal burden count of the spleen across the time points (Figure 6C). We observed minimal detection of fungal cells at 3 dpi within the spleen, as anticipated, given the spleen’s role for antigen presentation and immune response activation. A significant increase in fungal burden by 10 dpi was defined, which we anticipate is due to the presence of fungal cells circulating within the blood and filtering through the spleen; notably, several mice had minimal to non-detectable fungal levels at this time. Finally, a significant increase in fungal burden by 21 dpi was observed with all mice displaying high loads of *C. neoformans* within the splenic samples, which we attribute to increased dissemination of fungal cells in the blood and colonization of the spleen. Together, these fungal burden data support the proteome distinctions across the time points.

**Figure 6:**
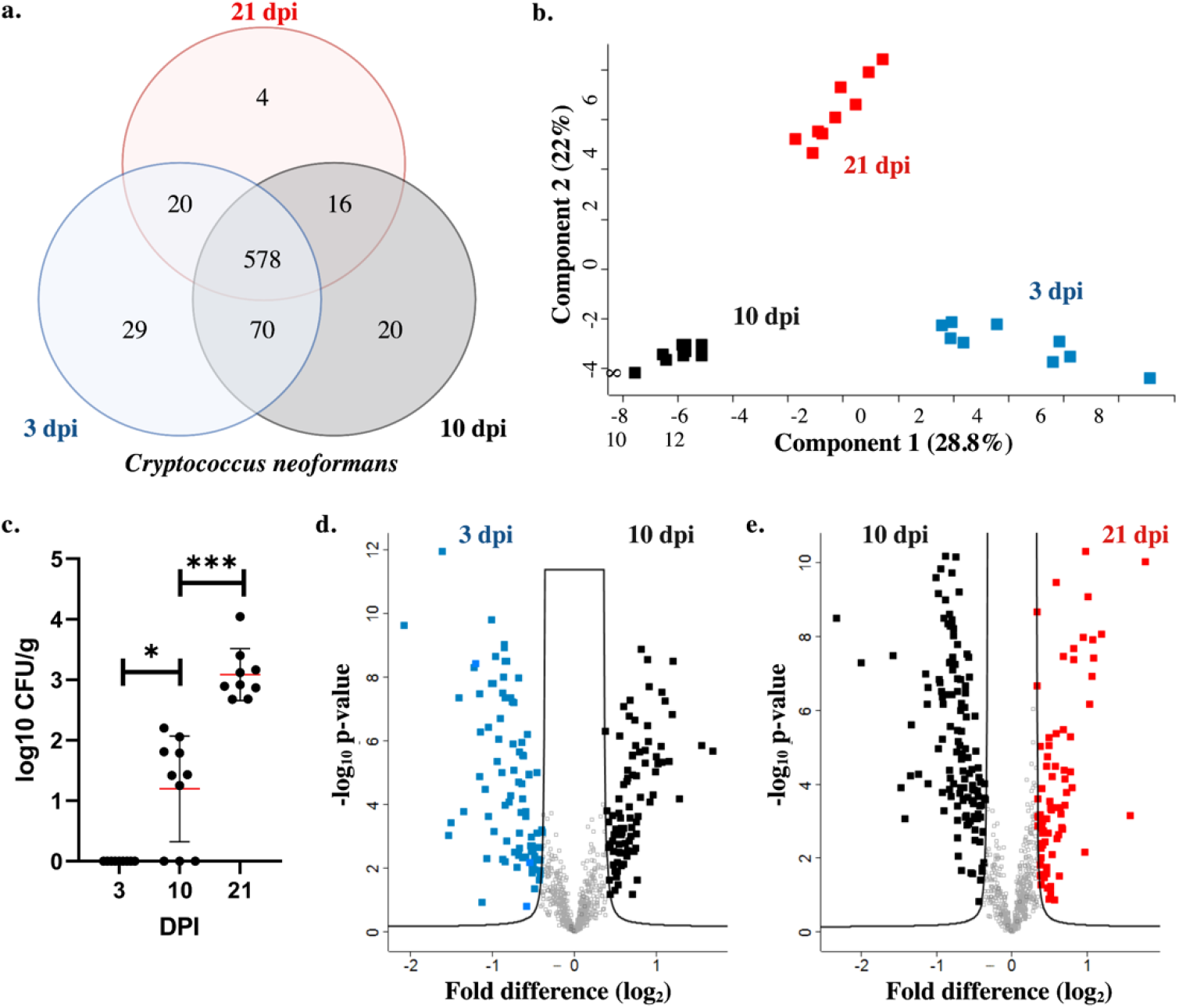
Pathogen-specific drivers of cryptococcal infection. A) Temporal comparison of identified proteins. A core proteome of 578 fungal proteins was observed with 29 proteins unique to 3 dpi, 20 proteins unique to 10 dpi, and 4 proteins unique to 21 dpi. B) Principal Component Analysis of fungal proteins from infected samples over time. C) Colony forming uni counts (normalized by splenic mass) for *C. neoformans* at 3, 10, and 21 dpi. Statistical comparison by Welch’s t-test, *p-value < 0.05, ***p-value < 0.001. D) Volcano plot for significantly different fungal proteins with higher production at 3 dpi (blue) vs. 10 dpi (black). E) Volcano plot for significantly different proteins with higher production at 10 dpi (black) vs. 21 dpi (red). F) 1D annotation enrichment based on Gene Ontology Biological Processes for comparisons of 3 dpi vs. 10 dpi and 10 dpi vs. 21 dpi. For volcano plots, Welch’s t-test p-value < 0.05, FDR = 0.05, S_0_ = 1.

Next, to assess significant changes in the fungal proteome during the course of infection, we performed statistical analyses on time point comparisons. At 3 dpi, we observed a significant increase in production of 88 fungal proteins compared to a significant increase in production of 87 proteins at 10 dpi (Figure 6D). A comparison of the later time points revealed 127 proteins with significant increases in abundance at 10 dpi, and 79 proteins with significantly increased abundance at 21 dpi (Figure 6E). For example, at 3 dpi, an adaptor protein associated with vesicle transport (CNAG_03928) showed the largest increase in abundance compared to 10 dpi when we observed an increase in abundance of polysaccharide capsule-associated proteins (mannose-1-phosphoate guanylyltransferase; mannose-6-phosphate isomerase). Conversely, at 21 dpi, we detected protein transport (Sec31), protein degradation (ClpB protease), and signaling cascades associated with fungal virulence (cAMP-dependent protein kinase regulatory subunit). Moreover, significantly different proteins across the time points included several virulence-associated fungal proteins, which we assess further in the following section. Together, these data support pathogen-specific responses throughout infection and highlight the influence of fungal burden on proteome remodeling at the later time point.

### 3.6. Predicting novel virulence factor functions and putative fungal biomarkers based on mapped patterns of production

To investigate the production of fungal proteins and predicted putative functionality as biomarkers, we focused on previously characterized *C. neoformans* virulence-associated proteins with distinct abundance profiles and fungal proteins with comparable patterns of production based on Pearson correlation (Table 2). For example, upon quantitative proteomic profiling of the spleen, we observed production of the cAMP-dependent protein kinase regulatory subunit, Pkr1(CNAG_00570), which has extensively well-defined roles in modulating fungal virulence and regulation of the cAMP protein kinase A pathway ^45, 46^. Production of the regulatory subunit was elevated at 3 dpi with a steady decline across the remaining time points (Figure 7A).

**Table 2:** Putative fungal protein biomarkers based on mapped patterns of production.

| Parent Trend Protein | Protein Name | CNAG ID | Protein ID | Keywords |
| --- | --- | --- | --- | --- |
| **CNAG_00570** | **cAMP-dependent protein kinase regulatory subunit** | **CNAG_00570** | **J9VH50** | **Kinase, Transferase** |
|  | Carbonic anhydrase | CNAG_05144 | J9VPA7 | Lyase |
|  | Mitochondrial import inner membrane translocase subunit | CNAG_05844 | J9VPH9 | Chaperone |
|  | Importin subunit beta-1 | CNAG_00988 | J9VRB1 | Protein transport |
|  | Large subunit ribosomal protein L7/L12 | CNAG_05980 | J9VSB4 | Ribosomal protein |
|  | S-methyl-5'-thioadenosine phosphorylase | CNAG_00165 | J9VDD8 | Glycosyltransferase |
| **CNAG_03007** | **CipC protein** | **CNAG_03007** | **J9VPP8** |  |
|  | Eukaryotic translation initiation factor 3 subunit D | CNAG_03641 | J9VIB2 | Initiation factor, Protein biosynthesis |
|  | Cytoskeleton assembly control | CNAG_02237 | J9VQ91 | Actin-binding |
|  | Non-histone chromosomal protein 6 | CNAG_06544 | J9VT78 | DNA-binding |
|  | Integral membrane protein | CNAG_01761 | J9VZA0 | Membrane |
|  | FK506-binding protein 1 | CNAG_03682 | O94746 | Isomerase |
| **CNAG_02189** | **Alpha-amylase** | **CNAG_02189** | **J9VQD3** | **Hydrolase** |
|  | Large subunit ribosomal protein L28e | CNAG_04068 | J9VHC1 | Ribonucleoprotein |
|  | Arginosuccinase | CNAG_02825 | J9VHI3 | Lyase |
|  | BZIP domain-containing protein | CNAG_07560 | J9VKV9 |  |
|  | Nucleolin | CNAG_07362 | J9VLN9 | RNA-binding |
|  | Sterol 24-C-methyltransferase | CNAG_03819 | J9VNP9 | Methyltransferase |
| **CNAG_05540** | **Urease** | **CNAG_05540** | **O13465** | **Hydrolase** |
|  | Uncharacterized protein | CNAG_00995 | T2BME6 |  |
|  | Histone H1 | CNAG_04168 | J9VRP1 | DNA-binding |
|  | Large subunit ribosomal protein L23 | CNAG_01976 | J9W225 | Ribonucleoprotein |
|  | Sulfide:quinone oxidoreductase | CNAG_01981 | J9W230 |  |
|  | Cytochrome c oxidase assembly protein subunit 17 | CNAG_05573 | J9W447 | Chaperone |

**Figure 7:**
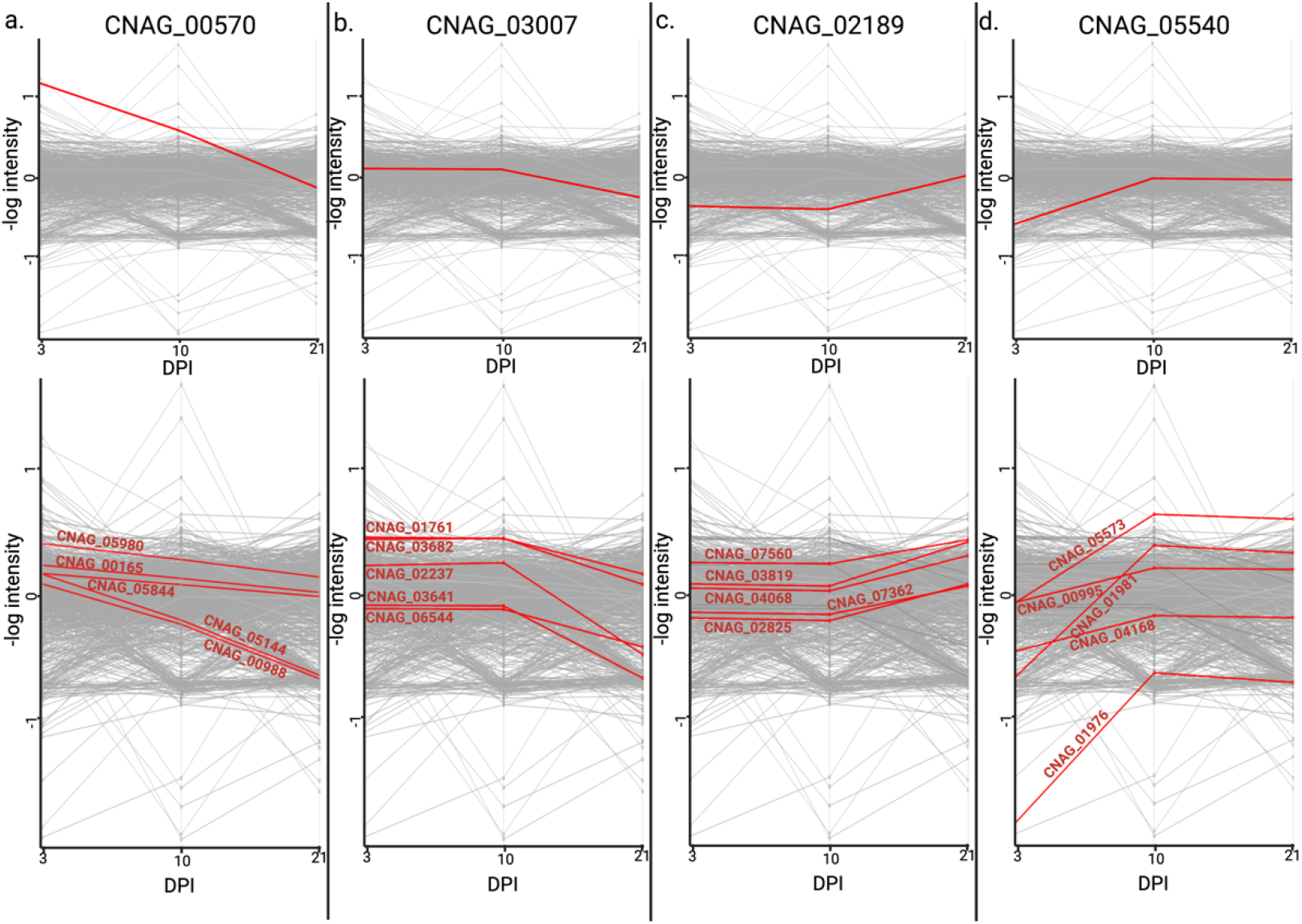
Mapped temporal production for virulence-associated fungal proteins. A) CNAG_00570 fungal protein mapped abundance trend at 3, 10, and 21 dpi. Top five fungal proteins with comparable mapped protein production based on Pearson correlation. B) CNAG_03007 fungal protein mapped abundance trend at 3, 10, and 21 dpi. Top five fungal proteins with comparable mapped protein production based on Pearson correlation. C) CNAG_02189 fungal protein mapped abundance trend at 3, 10, and 21 dpi. Top five fungal proteins with comparable mapped protein production based on Pearson correlation. D) A) CNAG_05540 fungal protein mapped abundance trend at 3, 10, and 21 dpi. Top five fungal proteins with comparable mapped protein production based on Pearson correlation.

Proteins mapping with comparable production profiles include, lyase (CNAG_05144), chaperone (CNAG_05844), transporter (CNAG_00988), ribosomal (CNAG_05980), and glycosyltransferase (CNAG_00165). We also detected CipC (CNAG_03007) with known roles in *C. neoformans* virulence within a murine model of infection^47^. Here, CipC had stable abundance at 3 and 10 dpi followed by reduced production by 21 dpi (Figure 7B). We observed comparably mapped proteins, including an initiation factor (CNAG_03641), actin-binding (CNAG_02237), DNA-binding (CNAG_06544), membrane (CNAG_01761), and isomerase (CNAG03682). Next, we focused on the production of ⍰-amylase (CNAG_02189) with previously defined roles in *C. neoformans* virulence and demonstration as a biomarker for cryptococcal infection from the blood of a murine model^48–50^. ⍰-Amylase showed consistent levels of protein production at 3 and 10 dpi followed by an increase in production at 21 dpi (Figure 7C). Proteins mapped with similar abundance profiles include ribonucleoprotein (CNAG_04068), lyase (CNAG_2825), RNA-binding (CNAG_07362), methyltransferase (CNAG_03819), and uncharacterized (CNAG_07560). Lastly, we identified urease (CNAG_05540), a hydrolase with importance for *C. neoformans* virulence, including pathogenesis within a macrophage environment and invasion of the central nervous system^51–54^. Urease was low at 3 dpi but increased in abundance at 10 and 21 dpi; with timing correlating with its role in tissue invasion for fungal dissemination throughout the host during infection (Figure 7D). We observed proteins associated with DNA-binding (CNAG_04168), ribonucleoprotein (CNAG_01976), oxidoreductase (CNAG_01981), chaperone (CNAG_05573), and uncharacterized (CNAG_00995) with similar production profiles. Based on protein production patterns, our data support potential novel roles for fungal proteins in virulence and suggest the discovery of time-dependent biomarkers with defined roles in *C. neoformans* virulence.

## 4. Discussion

In this study, we develop an innovative pipeline for proteomic profiling of infected murine tissue to uncover new biological drivers of cryptococcosis. Our dual perspective approach to applications of mass spectrometry-based proteomics for infectious disease research defines the temporal host response to *C. neoformans* infection and observes fungi-specific immune-associated proteins with altered abundance. We highlight pathogenic drivers of disease within the spleen, observing an initial splenic response to *C. neoformans* through antigen presentation and activation of innate and adaptive immune cells, followed by substantial remodeling of both biological systems in defense (host) and virulence (pathogen). These states are followed by fungal colonization within the spleen, providing a connection to dissemination of *C. neoformans* via the blood as the disease progresses; this information is correlated with quantification of fungal burden loads. Finally, we map patterns of host and pathogen protein production to propose novel functions for proteins based on abundance profiles and propose the exploration of putative biomarker signatures from complementary biological systems for monitoring the presence and progression of cryptococcal disease.

The spleen is an important secondary lymphoid organ, which serves as a reservoir for naïve B- and T-cells that become activated upon interaction with antigens presented by innate immune cells (e.g., macrophages, dendritic cells)^19^. Importantly, the antigen presenting cells, as well as pathogen cells and activated immune cells circulate throughout the body via the blood. This provides an important connection for the discovery of spleen-associated putative biomarkers that may circulate via the blood. Here, we do not detect *C. neoformans* cells within the spleen at 3 dpi but we do observe remodeling of the host proteome and detection of fungal proteins within the spleen, supporting antigen presentation from cells migrating from the initial site of infection (lungs) to activate an immune response (spleen). As anticipated from the host perspective, we identified multiple lectin-like receptors that play key roles in recognition of fungal pattern associated molecular patterns to initiate a response from innate immune cells^55, 56^. The abundance of these receptors is detected at 3 dpi and then increases significantly between 10 and 21 dpi, suggesting early and prolonged activation to mount an effective immune response, correlating with early detection of cryptococcal proteins within the spleen.

From the pathogen perspective, we achieve a depth of proteome profiling not previously reported for *C. neoformans* within the spleen. Our approach enables temporal identification of known and candidate novel *C. neoformans* virulence factors, including Pkr1, CipC, urease, and ⍰-amylase with known roles in fungal virulence, as well as and demonstrated mapped production patterns to other fungal proteins with known or putative roles in virulence. The timing of protein induction represents an opportunity to define a unique signature of disease based on presence and abundance of these proteins upon *C. neoformans* infection. Proteins with mapped production profiles matching these virulence factors, such as chaperones and lyase, have known roles in cryptococcal virulence, providing validity to the comparison. Molecular chaperones, such as heat shock proteins, are well characterized chaperones with key roles in *C. neoformans* stress tolerance, thermotolerance, and virulence^57, 58^. Lyases are also associated with fungal virulence and represent potential anti-virulence targets (i.e., disrupt the production and assembly of virulence factors and interfere with virulence factor function)^59–62^. Additionally, the FK506-binding protein 1 isomerase (mapped to CipC) has defined roles as a target for rapamycin antifungals^63^ and two proteins mapped to urease (chaperone and oxidoreductase) are important for fungal defense against reactive oxygen species produced by the host during infection^64, 65^. Other virulence associated fungal proteins identified by our analysis include an ABC transporter that confers fluconazole resistance^66^ (a mainstay antifungal drug used to treat cryptococcosis), AFR1, with a significant increase in production at 3 dpi. This protein also promotes resistance of *C. neoformans* to microglia-mediated antifungal activity through delayed maturation of the phagosome^67^. Together, our approach defines novel and known mechanisms of fungal virulence, suggesting putative biomarker signatures to track disease over time, as well as potential new targets for anti-virulence intervention strategies.

Over the past two decades mass spectrometry-based proteomics has increased in popularity, accessibility, and reproducibility due to technological and bioinformatic advances^68^. For infectious disease biology, applications of mass spectrometry as a diagnostic tool, known as proteotyping, have become routine, and although, such processes are cost effective, automated, and relatively rapid, limitations still exist^1, 69^. For example, distinction between species and strains, assessing non-culturable pathogens, efficacy towards fungal pathogens, and providing information about the severity of infection, limit the diagnostic power^2^. To overcome such challenges, high resolution mass spectrometry combined with bioinformatics and artificial intelligence are gaining in popularity^5^. Our group, and others, are pioneering dual perspective proteomics to define global proteome changes upon infection across multiple biological systems^44, 70^. These datasets provide new insight into the host mechanisms responsible for protection from infection and unparalleled insight into pathogenic responses.

## 5. Conclusion

Here, our application of temporal state-of-the-art mass spectrometry-based proteomics of the spleen following cryptococcal infection reveals anticipated and novel responses of the host and known and candidate virulence-associated factors of the pathogen. We propose proteins across the biological systems as putative protein biomarkers that can track disease progression and severity over time and aim to provide unique patterns of response for the design of a biomarker signature unique to *C. neoformans*. Such approaches of protein biomarker signatures are being developed for neurological disorders^71^, aging^72^, and cancer^73–75^, as well as recently advanced for COVID-19^76^ applications; however, contributions to infectious diseases, particularly for fungal pathogens, have been severely limited^77^. Here, we develop a pipeline for the comprehensive temporal proteomic profiling of *C. neoformans* infection using a murine model with spatial consideration confined to the spleen; however, our discoveries support new biological insight into mechanisms driving cryptococcosis and propose a strategy to explore tissue-specific biomarkers for effective and efficient diagnosis, monitoring, and treatment of cryptococcosis.

## Author Contributions

B.M. & J.G.-M. conceptualized and designed the study. B.M. & F.R.-D. performed experiments and data analysis and designed and developed figures. F.R.-D. measured samples on the mass spectrometer and performed the preliminary raw file processing. B.M. & J.G.-M. generated figures and wrote the first draft of the manuscript. B.M., F.R.-D., A.D. & J.G.-M. edited the manuscript and wrote the final version. All authors have read and approve the submitted manuscript.

## Acknowledgments

The authors thank Kiyan Kheradvar for technical assistance with plating samples and counting colonies from the murine model experiments and members of the Geddes-McAlister lab for their informative and constructive feedback on project design and manuscript preparation.

## Data Availability

The RAW and affiliated files were deposited into the publicly available PRIDE partner database for the ProteomeXchange consortium with the data set identifier: PXD040868

Reviewer login

Password: bkVB6DY6

